# Anti-vitronectin single-chain variable fragment targeting the hemopexin-like domain decreases age-related macular degeneration indicators in ex vivo models of the retinal pigment epithelium

**DOI:** 10.1101/2025.08.18.670968

**Authors:** Jun W. Kim, Susanne Meyer, Ben Lopez, Mohamed Faynus, Lincoln V. Johnson, Houman Hemmati, Dennis O. Clegg, Hyunsuk Min

## Abstract

Vitronectin is a prominent constituent of the extracelluar matrix (ECM) that plays a key role in inflammation by regulating cell adhesion, migration, and complement activation. Together with other inflammatory biomarkers, lipids, and hydroxyapatite, an ECM component, vitronectin, accumulates in abnormal deposits and drusen associated with age-related macular degeneration (AMD). Using an engineered polypeptide (VnHX) containing the hexmopexin-like domain (HX) of vitronectin, we found that the HX domain directly promotes hydroxyapatite (HAP) accumulation. Next, we screened humanized single-chain variable fragments (scFvs) against vitronectin that inhibit the interaction between VnHX and HAP, leveraging the binding of VnHX to HAP. We then tested these scFvs for their ability to block Vn-containg deposits secreted by stem cell-derived retinal pigment epithelium (RPE). In this in vitro drusen model, treatment with the anti-vitronectin antibodies decreased vitronectin accumulation. Furthermore, the antibody treatment led to decreased accumulation of C5b-9 and clusterin, indicating alterations in the complement pathway and cellular stress. These results support that vitronectin has a functional role drusen accumulation and possibly in AMD progression. Vitronectin would be a novel, promising therapeutic target for AMD.

## INTRODUCTION

Age-related macular degeneration (AMD) is a predominant cause of vision loss in individuals aged 55 and above, affecting approximately 8.7% of the population between 45 and 85 years old^1^. This irreversible central vision loss significantly hampers daily activities, emphasizing the urgent need for effective detection and treatment. While current approaches focus on managing the disease through supplements and diet, their efficacy is limited, especially in advanced stages. The multifactorial nature of AMD, influenced by genetic susceptibility, age, lipid metabolism, and immune dysregulation, poses a challenge to fully understanding its mechanism^2^. Although recent advancements have introduced therapies targeting specific pathways, such as anti-angiogenic treatments and complement pathway inhibitors^3-5,^, their variable effectiveness and inability to prevent long-term vision loss underscore the existing gaps in AMD management.

AMD primarily affects the photoreceptors, retinal pigment epithelium (RPE), Bruch’s membrane (BrM) and the choroid within the macula. The impairment of these layers results from dysregulated lipid metabolism and immune system functions, which are influenced by both aging and environmental factors. Perturbations in RPE’s metabolic pathways lead to the accumulation of lipids effluxed from the RPE, such as lipofuscin, creating acellular debris known as drusenoid deposits within the BrM. These deposits, composed of oxidized lipids, lipoproteins, and immunogenic complement components, are the most recognized risk factor for AMD, and their growth prompts the detachment of RPE cells from their membrane, facilitating their migration into the neurosensory retina^6^. This process corresponds with the advancement of the disease, culminating in the loss of affected layers in the advanced stages of dry AMD or the invasion of new vessels and fibrosis in neovascular AMD (wet AMD). Consequently, or concomitantly with these processes, the inflammatory system exacerbates tissue damage. Hence, exploring the extracellular aggregates linked to the structural and cellular changes in AMD could unveil novel therapeutic strategies for the disease.

Vitronectin is a prominent constituent of the extracellular matrix (ECM) that plays a key role in angiogenesis and inflammation by facilitating cell adhesion, migration, and complement activation^7,8^. It is also associated with abnormal deposits in various diseases, including AMD and atherosclerosis^9,10^. In human and porcine cell cultures, it has been confirmed that vitronectin, forms an even coat on these deposits^11,12^. A genetic variant of vitronectin has also been associated with AMD^13^. Structurally, vitronectin has the central and the C-terminal domains (HX1 to HX4) homologous to the hemopexin (HX) family^14^. Recent studies show that the HX domains of vitronectin (VnHX) binds to hydroxyapatite (HAP) microspheres^15^. NMR results show an alignment between the rim-Asp in VnHX align and HAP calcium ion positions, facilitating the exchange of ionic calcium between vitronectin and HAP. HAP is also a major component of drusenoid deposits known to accumulate with lipid^11^.

Despite the previous studies associating vitronectin with AMD, the functional involvement of vitronectin in AMD has been unclear. In this study, we employed an ex vivo model using stem-cell derived RPE, which secrete drusen-like deposits in culture. We showed that in ex vivo models of RPE, laminin coating as a substrate and the treatment with human serum promoted vitronectin accumulation, indicating the model’s utility for examining vitronectin’s role in drusen accumulation Then, by taking advantage of the binding interaction between VnHX and HAP we identified humanized single-chain variable fragments (scFvs) that inhibit the interaction. The treatment with the anti-vitronectin antibodies decreased the accumulation of vitronectin in the ex vivo RPE model. Moreover, the antibodies decreased the levels of the terminal complement complex C5b-9 and the oxidative stress marker, clusterin. The differentiation state of the cells was unaffected, as shown by unaltered expression levels of ZO-1 and RPE65. These results indicate the functional role of vitronectin in drusen accumulation.

## RESULTS

### Laminin promotes vitronectin accumulation in ex vivo RPE

Evidence from prior studies indicate that dysregulation of RPE function in AMD leads to the accumulation of metabolic deposits followed by retinal cell state alteration and death (Figure 1a). BrM serves as a basement membrane to RPE, providing cellular anchorage, acting as a barrier and a filter, and stabilizing the overall structure of the retinal tissue^16^. Because BrM is functionally involved in RPE pathologies such as AMD, ex vivo models frequently include BrM-mimicking substrates. Matrigel, a heterogeneous mouse tumor-derived mixture, is often used to supply ECM components in both two-dimensional (2D) and three-dimensional (3D) settings, although it has non-specific physiological relevance^17^. One of the major components of BrM is laminin, to which RPE cells bind through integrins^18^. In addition to mechanical support, laminin has been shown to affect barrier status and promote differentiation of RPE^18^.

**Figure 1.**
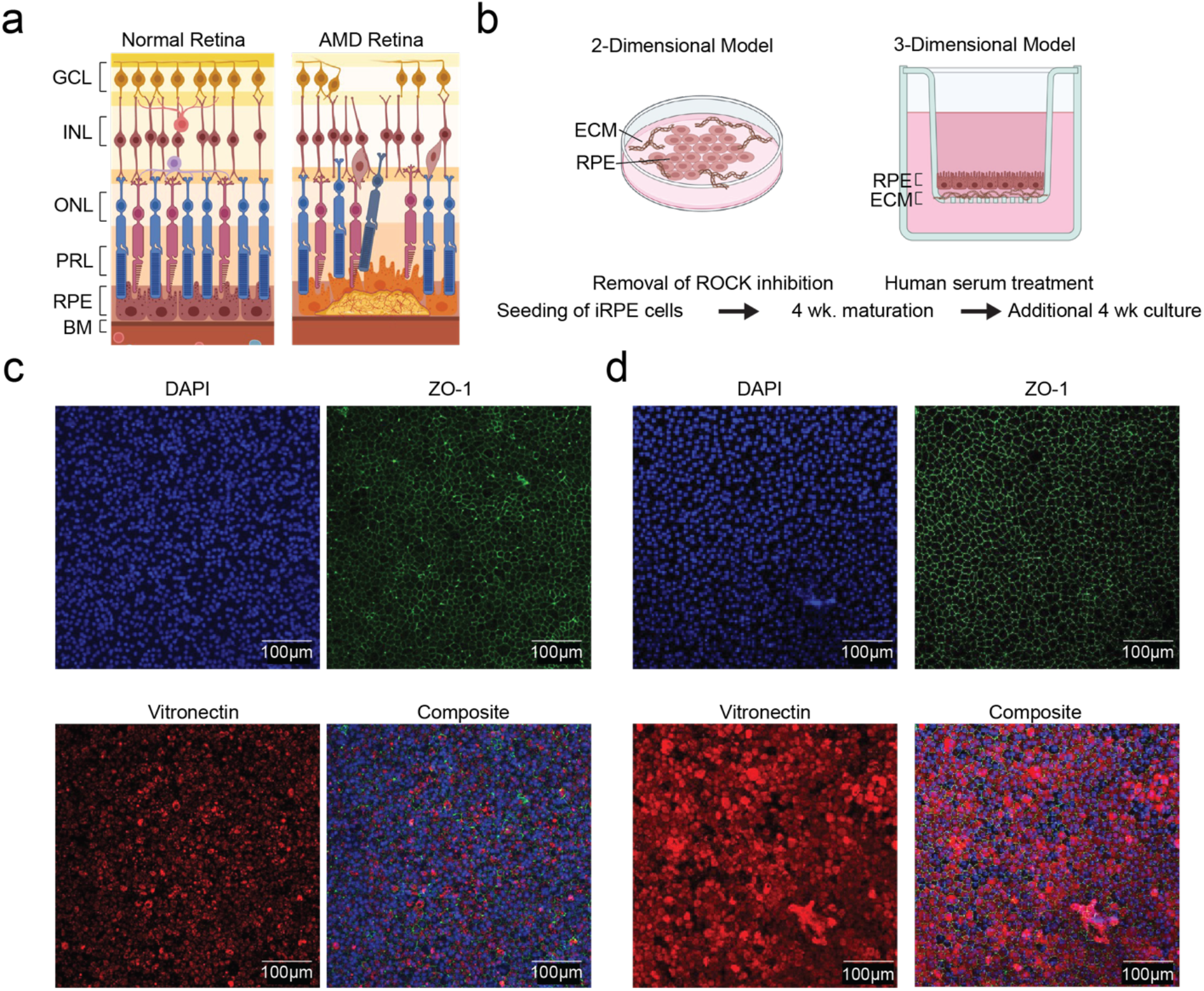
Ex vivo retinal pigment epithelium (RPE) models show laminin induced vitronectin accumulation, **a**. Schematic of retinal anatomy and age related macular degeneration (AMD) progression. AMD affects the photoreceptors, RPE, Bruch’s membrane and the choroid within the macula. These impairments are associated with dysregulated lipid metabolism and immune system functions. GCL: ganglion cell layer; INL: inner nuclear layer; ONL: outer nuclear layer; PRL: photoreceptor layer; RPE: retinal pigment epithelium; BM: Bruch’s membrane, **b**. Two-dimensional (2D) and three-dimensional (3D) ex vivo models of RPE on extracellular matrix (ECM). The 3D model uses a porous membrane that allows bidirectional nutriend diffusion. Two different ECMs (matrigel and laminin) were used to mimic the function of Bruch’s membrane. The models were matured for a total of 8 weeks before analysis, **c**. Differentiated RPE cells on a 2D model with matrigel, fluorescently labeled for nucleus (DAPI), ZO-1 (green), and vitronectin (red), **d**. Same as in **c**. but with laminin, **e**. Differentiated RPE cells on a 3D model with matrigel and laminin, fluorescently labeled for nucleus (DAPI), ZO-1 (green), ApoE (red), and vitronectin (yellow).

In both 2D and 3D ex-vivo human RPE models, cells were cultured for 4 weeks without ROCK inhibition to induce differentiation (Figure 1b)^19^. Both Matrigel and laminin supported RPE differentiation, as demonstrated by the cobblestone morphology and expression of ZO-1 and RPE65 (Supplementary Figure 1a). In the 3D model, minor autofluorescence was detected from the porous membrane, but a similar observation was noted (Supplementary Figure 1b). 3D projections of confocal images revealed both apical and basal APOE (Supplementary Figure 1c). Compared to the Matrigel-based model, RPE cells cultured on laminin showed higher accumulation of vitronectin (Figure 1c, d), reflecting the AMD phenotype^20^. Consequently, laminin was used as the primary laminin ECM support for both 2D and 3D models.

### Phage display screening of anti-vitronectin antibodies that prevent HAP accumulation

As a key factor in AMD progression, RPE cells synthesize and secrete lipoprotein components integral to lipid metabolism, uptake, and signaling^21^. RPE-secreted lipids contribute to both subretinal and basal deposits, which can lead to RPE cell detachment, migration, or death^22^. These deposits are composed of multiple components, among which HAP spherules serve as scaffolds for additional protein accumulation^11^. Vitronectin binding to HAP spherules has been demonstrated and forms the mechanistic link to AMD pathogenesis^15^.

Although laminin-induced vitronectin accumulation was evident in a three-month ex vivo culture, we did not detect HAP deposits, likely due to the relatively short culture duration. To investigate vitronectin’s functional role in HAP formation, we first confirmed the binding interaction between VnHX and HAP using a co-sedimentation assay, in which sequestration of supernatant VnHX by HAP microspheres was detected using PAGE analysis (Supplementary Figure 2a, b). To identify antibodies capable of blocking the vitronectin-HAP interaction, we performed phage display screening of single-chain variable fragments (scFv) targeting the HAP-binding region of VnHX (Figure. 2a). Briefly, a naïve human antibody library (∼5×10^10^ clones) was panned against an avi-tag-fused VnHX (avi-VnHX) immobilized on biotin-coated surface. HAP microparticles were added to mask the HAP-binding surface on VnHX, and only phages displaying scFvs that compete with HAP at this site were eluted and amplified. After four rounds of selection, seven unique scFv sequences were identified. Each of these clones was expressed with His-tag (HIS6x) for purification and subsequent binding analysis (Supplementary Figure 2c).

**Figure 2.**
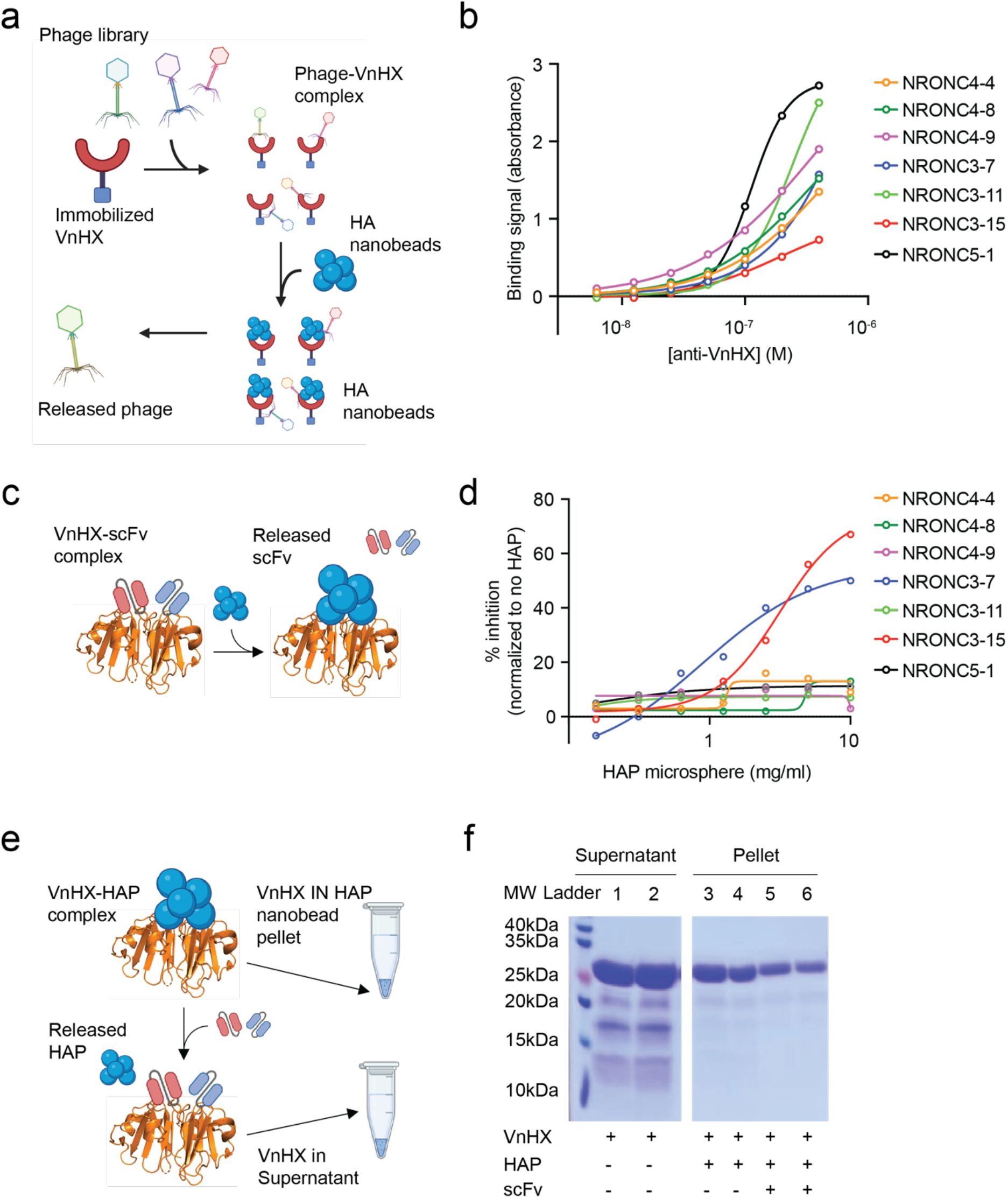
Discovery of neutralizing anti-VnHX antibodies (scFv). **a**. Schematic showing phage display screening workflow to discover neutralizing anti-VnHX antibodies, **b**. Binding curves of the solubly expressed top 7 antibodies isolated from the phage display screening in A. **c**. Schematic showing competition between antibodies and HAP for VnHX. Adding high concentration of HAP to VnHX bound to a neutralizing antibody is expected to release the antibody from VnHX. **d**. % inhibition of antibody binding to VnHX by HAP microspheres, **e**. Schematic of co-sedimentation assay showing that VnHX-HAP complex is expected in the HAP microsphere pellet, while VnHX-antibody complex is expected in the supernatant, f. Reduced PAGE analysis of co-sedimentation assay with Coomassie blue staining for VnHX with and without the neutralizing antibodies (60 pg/mL). NRONC3-7 is used for lane 5, and NRONC3-15 is used for lane 6.

An ELISA confirmed that all scFvs bound to immobilized VnHX with signals above background at submicromolar concentrations (Figure 2b). Next, we assessed their specificity for the HAP-binding domain via a competition assay using HAP nanobeads: the scFvs (5 μg/mL) were mixed with varying concentrations of the HAP microbeads (0.15-10 mg/mL) incubated with VnHX-coated wells; the fraction of scFvs that remained bound was quantified using ELISA (Figure 2c). Among the seven VnHX binders, two constructs, NRONC3-7 and NRONC3-15, robustly competed with HAP for the VnHX binding interface (Figure. 2d). We further analyzed these constructs in a pull-down assay (Figure 2e). In agreement with the ELISA results, adding the scFvs to a solution of VnHX mixed with HAP microbeads reduced the amount of VnHX retained on the HAP nanobeads, as shown by PAGE analysis (Figure 2f).

### Anti-vitronectin antibody treatment on ex-vivo RPE model decreases vitronectin accumulation and AMD-associated biomarkers

A short-term (48 h) exposure to 200 μg/mL of NRONC3-7 or NRONC3-15 did not show detectable changes in RPE morphology or pigmentation (Supplementary Figure 3). We therefore proceeded to investigate their capacity to modulate AMD-associated phenotypes in our ex vivo models. 2D and 3D RPE models were matured for four weeks and exposed to human serum, which elevated the AMD-associated complement marker C5b-9 and the inflammatory marker serum amyloid P component (APCS; Supplementary Figure 4a). The cultures were then incubated with 200 μg/mL of NRONC3-7 and NRONC3-15 for an additional four weeks, with fresh medium (containing the respective antibodies) replenished twice weekly to maintain consistent concentrations (Figure 3a).

**Figure 3.**
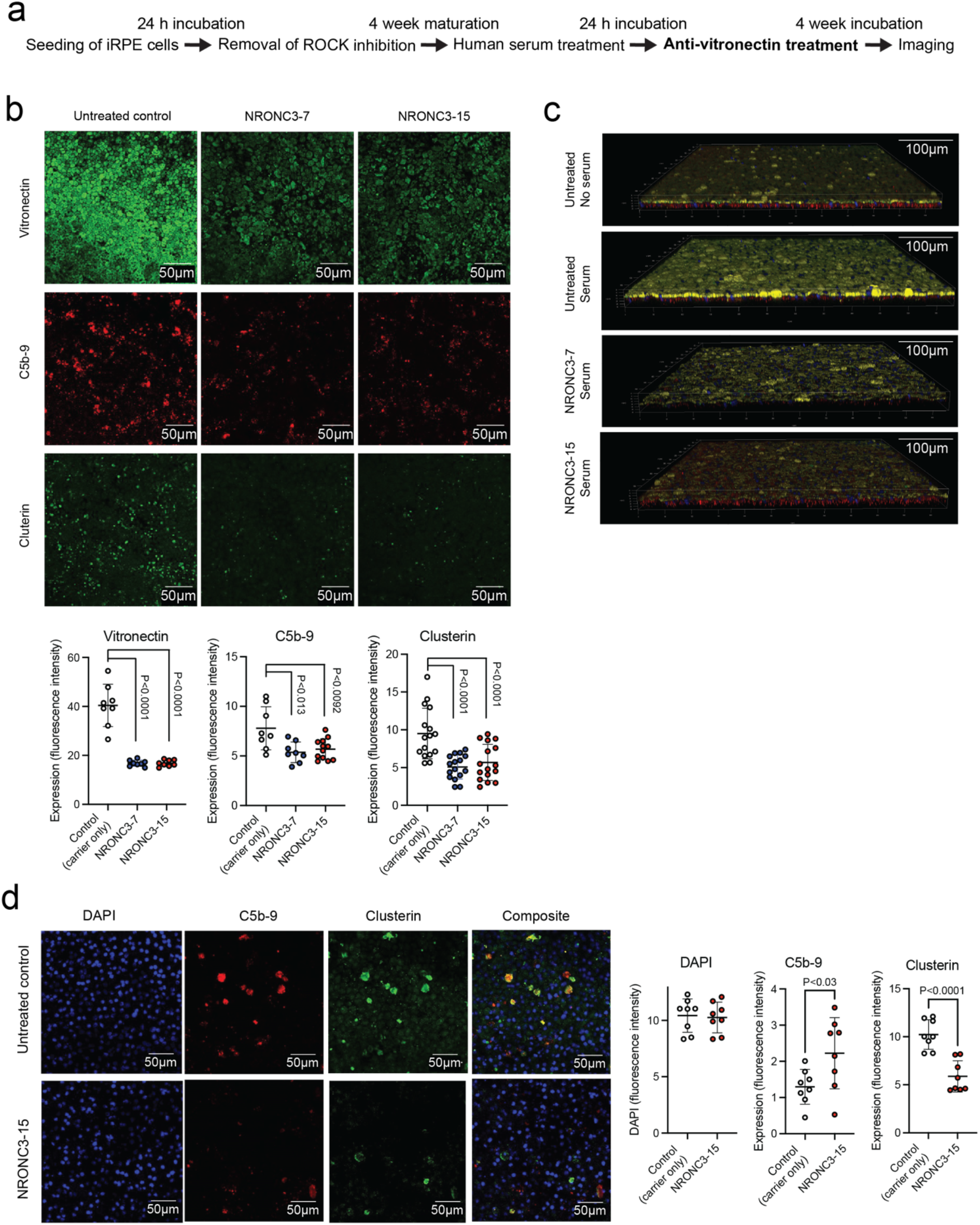
Anti-VnHX scFv decreases accumulation of vitronectin, C5b-9, and clusterin. **a**. RPE model preparation timeline. The samples were treated with the selected anti-VnHX scFvs (NRONC3-7 and NRONC3-15) for 4 weeks before imaging analysis, **b**. Control and scFv-treated 2D RPE models on laminin were fluorescently labeled for vitronectin (green), C5b-9 (red), and clusterin (green), **c**. Control and scFv-treated 3D RPE models on laminin were fluorecetnly labeled for nucleus (DAPI), ZO-1 (green), ApoE (red), and vitronectin (yellow), **d**. Same as in c. but stained for nucleus (DAPI), clusterin (green), and C5b-9 (red). For b and d, the four quadrants of a sample was analyzed separately to account for the curvatures of the membranes. At least two separate samples were used for analysis. One way analysis of variance (ANOVA) were performed followed by unpaired Student’s t-test for post hoc test.

In the 2D model, immunofluorescence imaging revealed that, compared to untreated controls, NRONC3-7 and NRONC3-15 reduced vitronectin, C5b-9, and clusterin, while leaving APCS unchanged (Figure 3b). In the 3D laminin-based model, human serum treatment led to uniformly distributed apical vitronectin accumulation (Figure 3c). Although vitronectin can localize to both apical and basal surfaces of RPE, its apical distribution in laminin-treated conditions resembles the reticular pseudodrusen pattern^23^. By contrast, Matrigel-based 3D models treated with human serum did not show increased vitronectin (as in the 2D model), indicating that serum exposure alone does not account for all vitronectin accumulation.

Consistent with findings in the 2D model, the laminin-based 3D model exhibited reduced vitronectin, clustering, and C5b-9 after NRONC3-7 or NRONC3-15 treatment (Figure 3d). Longer cultures led to bending of the membrane supports, producing curved planes that required further processing of the images (Supplementary Figure 4b). Overall, these results suggest that anti-vitronectin antibodies targeting the HAP-binding domain of vitronectin can mitigate AMD-associated pathology.

## DISCUSSION

Previous studies have demonstrated that hydroxyapatite (HAP) is a functionally important component of pathogenic lipid accumulations in various diseases and that vitronectin may mediate hydroxyapatite deposition. Our findings confirm that vitronectin indeed promotes HAP accumulation. To evaluate the potential for antibody-mediated intervention to inhibit this process, we employed phage display to screen for anti-vitronectin antibodies specifically tailored to disrupt vitronectin-HAP interactions. By exploiting the binding between VnHX and HAP, we first panned a 5×10^10^ humanized library against VnHX, then further selected clones based on their ability to compete with HAP for vitronectin binding. Two antibodies exhibited robust affinity for VnHX and were validated to interfere with HAP binding, indicating that antibodies targeting the VnHX domain can effectively mitigate vitronectin-mediated HAP accumulation.

In AMD, vitronectin-and HAP-rich drusenoid deposits are frequently observed associated with RPE. Various ex vivo RPE models exist-broadly categorized as two-dimensional (2D) or three-dimensional (3D). Although 2D models facilitate higher-throughput analyses and leverage well-established protocols, they lack the native structural environment. In contrast, 3D models often employ a porous support that partially recapitulates Bruch’s membrane (BrM), but typically require more ad hoc optimizations. Matrigel is a common substrate for RPE seeding bcause of its growth factors, whereas laminin is one of the most abundant ECM components of BrM. Although both 2D and 3D conditions yielded robust and comparable RPE differentiation in Matrigel and laminin, laminin in 3D notably induced greater apical vitronectin accumulation, highlighting the significance of the support layer for this phenotype.

We next evaluated the effect of anti-vitronectin treatment in both 2D and 3D ex vivo RPE models on laminin. While the exact functional relevance of drusen in humans requires further elucidation, multiple lines of evidence suggest that they contribute to AMD progression. Both HAP and vitronectin are key components of drusenoid deposits. Treatment with anti-vitronectin did not alter RPE markers, suggesting that the differentiation status of RPE cells remained intact, yet it did reduce vitronectin accumulation. The recent approval of a complement system inhibitor points to immune dysregulation as a viable therapeutic target in AMD, and C5B-9, a critical complement pathway marker, was also reduced alongside vitronectin. Clusterin, another drusen-associated component with roles in cellular homeostasis and stress, likewise decreased following anti-vitronectin treatment, potentially indicating a less stressed cellular environment, though further validation is needed.

In summary, we identified and characterized anti-VnHX antibodies that inhibits HAP binding and attenuates the accumulation of multiple drusenoid molecules implicated in AMD progression. These findings suggest that the HX domain of vitronectin constitutes a promising drug target for AMD through multiple pathways. Beyond AMD, the presence of vitronectin and HAP is also noted in other pathological settings, including cardiovascular disease, further underscoring the potential therapeutic relevance of disrupting vitronectin-mediated HAP interactions.

## MATERIALS AND METHODS

### iRPE cell culture

iCell Retinal Pigment Epithelial (RPE) cells (Catalog #C1047) were obtained from FUJIFILM Cellular Dynamics, Inc. Each vial contained 1.6 × 10^6^ RPE cells derived from human induced pluripotent stem cells (iPSCs). These cells represent a highly pure population of RPEs originating from the blood of a healthy Caucasian male donor.

On Day 0, one vial of iCell RPE cells (1.6 × 10^6 cells) was thawed and resuspended in X-Vivo-10 medium (Lonza, Inc.) with 10 µM Rock inhibitor (Y-27632, Tocris). Cells were plated into a single well of a six-well plate pre-coated with Matrigel (Corning, Inc.). Cultures were maintained at 37°C and 5% CO_2_, and fed twice weekly with fresh X-Vivo-10 medium with 10 µM Rock inhibitor. On Day 15, confluent cells were enzymatically lifted from the six-well plate using TrypLE (Thermo Fisher Scientific). The cell suspension was centrifuged at 300 × g for 5 minutes, and the pellet was resuspended in X-Vivo-10 medium with 10 µM Rock inhibitor. Cells were re-plated into three Matrigel-coated 25 cm^2^ culture flasks. Cultures were once again maintained at 37°C and 5% CO_2_, with twice-weekly medium changes. On Day 18, RPE cells were subcultured again. Cells were detached with TrypLE, pelleted by centrifugation, and resuspended in X-Vivo-10 medium with 10 µM Rock inhibitor. The resulting suspension was evenly distributed into four Matrigel-coated 75 cm^2^ flasks. Cells continued to receive biweekly medium changes under the same incubation conditions. Phase-contrast images were recorded to assess RPE morphology and confluence. On Day 21, confluent RPE cultures were observed and imaged before freezing. Cells were harvested with TrypLE, pelleted, and resuspended in X-Vivo-10 medium for counting. Following a final spin, the cell pellet was gently resuspended in ice-cold CryoStor CS10 freeze medium (STEMCELL Technologies) at a density of 2 × 10^6^ cells per mL. Aliquots of 1 mL per vial were immediately placed into 21 cryovials (each vial containing 2 × 10^6^ cells). The cryovials were transferred to a Mr. Frosty freezing container and placed at −80°C for 24 hours. The vials were then moved to a liquid nitrogen tank for long-term storage.

### 2D RPE model

A vial of passage 4 (P4) RPE cells (2 × 106 cells per vial) was thawed according to the manufacturer’s instructions and plated directly onto either (i) laminin- or Matrigel-coated 96-well Ultra Microplates (Revvity, black, cell culture-treated, ultra-clear bottom; Fisher Scientific Cat. #50-209-936) or (ii) laminin- or Matrigel-coated cell culture inserts in a 24-well plate. Cells were maintained in X-Vivo-10 medium (Lonza), with medium changes performed twice weekly. After 4 weeks of maturation, cells were exposed to complement-competent serum to induce complement-mediated effects. The following day, cultures were treated with the candidate therapeutic antibody (0.5 mg/mL) or an appropriate control. Cells continued to receive medium changes twice weekly, with the antibody (or control) diluted in X-Vivo-10 medium.

After 4 weeks of the treatment period, cells were washed once with PBS and fixed in 4% paraformaldehyde (PFA), followed by another PBS wash. To reduce nonspecific binding, samples were blocked with block buffer (4% normal goat serum and 0.1% Triton x-100 in PBS). Primary antibodies were applied and incubated overnight at 4°C. The following day, samples were washed in PBS and incubated for 30 minutes with fluorescence-conjugated secondary antibodies and Hoechst nuclear stain. Finally, cells were washed with PBS and prepared for confocal microscopy analysis.

### 3D RPE model

RPE cells were seeded onto Millicell Standing Cell Culture Inserts (MilliporeSigma, Cat. #PIH01250) placed in Falcon 24-well plates (Fisher Scientific, Cat. #08-772-1), following the same culture conditions as the two-dimensional (2D) model. Cells were maintained as previously described, with regular medium exchanges.

### Immunofluorescence Preparation and Imaging

To prepare insert-grown RPE monolayers for imaging, culture medium was aspirated from both the inserts and the wells. Cells were briefly washed with PBS, fixed with 4% paraformaldehyde (PFA), and then rinsed again with PBS. The membrane supporting the fixed RPE cells was excised from the insert using a sterile scalpel and transferred to a clean 24-well plate. Non-specific binding was blocked by incubating the membranes with block buffer for 30 minutes at room temperature, followed by overnight incubation at 4°C with the relevant primary antibodies. After three washes in PBS, the membranes were incubated for 30 minutes with fluorescently labeled secondary antibodies and Hoechst nuclear stain, rinsed again in PBS, and mounted onto glass slides with ProLong Gold Antifade reagent (Thermo Fisher Scientific) under coverslips. Slides were allowed to cure before confocal microscopy.

### Co-sedimentation assay of VnHX and HAP nanospheres

Co-sedimentation assay was conducted according to the description from a prior study. Briefly, HAP microbeads (2.5 μm diameter, 100 m2/g surface area; Sigma-Aldrich catalog no. 900195) and SiO2 gel (60 nm diameter; Sigma-Aldrich catalog no. 288594) were washed three times in buffer M1 (20 mM MES, pH 6.5, 300 mM NaCl) by suspending in 100 mg of particules in 1 mL of buffer followed by centrifugation at 5,000 × g for 5 min in 4 °C. To assess VnHX sedimentation, 10 mg of HAP microbeads or SiO_2_ gel was gently mixed with 200 μL of 10 μM VnHX (kindly provided by Kyungsoo Shin and Francesca M. Marassi at Medical College of Wisconsin^14^) in buffer M1. Each mixture was centrifuged (5,000 × g, 5 min, 4 °C) to separate the solid fraction from the supernatant; the pellet was then washed with 200 μL of fresh buffer and recovered by an additional centrifugation step. Protein in each fraction were analyzed by Coomassie-stained SDS-PAGE.

### Phage display screening

Two proprietary naïve human scFv phage display libraries (∼5×10^10^ unique clones each) were obtained from NeoClone and ProteoGenix. Briefly, VL from Kappa and Lambda light chains were processed separately to reduce construction bias, and in-frame sequence frequencies were confirmed to exceed 95%. Two rounds of library screening against VnHX were performed to select an initial pool of binders. In each round, avi-tagged VnHX (avi-VnHX) was immobilized on biotin-coated tubes, incubated with the phage library, washed with PBS, and eluted using glycine-HCl. The eluted phages were propagated in E. coli train TG1, precipitated with PEG/NaCl, and resuspended for subsequent steps.

To further enrich for VnHX-neutralizing binders, VnHX was first incubated with hydroxyapatite microbeads (HAP; 2.5 μm diameter, 100 m^2^/g surface area; Sigma-Aldrich Cat. No. 900195) to block epitopes outside the HX domain. The previously enriched library was then exposed to this VnHX–HAP complex. Next, the filtered pool was incubated with avi-VnHX and competitively eluted with HAP microbeads. Phages recovered from this step were again infected into E. coli TG1, plated for counting, and used to calculate plaque-forming units (pfu). For an initial assessment of VnHX binding in the screened phage pool, a polyclonal phage ELISA was performed on plates coated with avi-VnHX (50 μg/mL). After washing and blocking, approximately 2 × 10^12^ amplified phages were added and allowed to bind. Following extensive washing, an anti-phage HRP-conjugated antibody was added. After another washing, TMB substrate was added, and then the reaction was stopped with HCl. Absorbance was measured at 450 nm. The same procedure was used for monoclonal phage ELISA to confirm individual binders to VnHX.

### Anti-vitronectin single-chain variable fragment production

The identified anti-vitronectin scFvs were expressed in mammalian cells for scale up production. Briefly, endotoxin-free expression vectors were used to transiently transfect 1-L culture of proprietary XtenCHO cells (ProteoGenix) in animal product-free medium Culture medium was collected when viability dropped <50% (14 days post-transfection). Recombinant antibodies were purified on a nickel resin using a standard method. Briefly, media containing the expressed scFv were clarified by 0.22 µm filtration. After the pH is adjusted to 7.5, the media was applied to nickel column to allow the His-tagged protein to bind to the nickel resin. The loaded column was washed and eluted by imidazole shift. The pool of fractions of interest was buffered exchanged with PBS. The final samples were qualitatively analyzed by PAGE and quantitatively by UV280 (>95% purity).

### Fluorescence imaging

Confocal fluorescence images were captured as .lif files on a Leica SP8 Resonant Scanning Confocal Microscope at 10x and 20x magnifications using Leica’s LASX software. LASX software was also used for generating 3D image files. .lif files were processed in ImageJ (Fiji) for preparation of 2D images.

## ACKNOWLEDGEMENTS

We would like to acknowledge the contributions of the members of Neuroptika. We also thank the members of the Clegg lab and the UCSB Stem Cell Center for helpful suggestions and Kyungsoo Shin and Francesca M. Marassi at Medical College of Wisconsin for providing VnHX.

## FIGURES

**Supplementary Figure 1.**
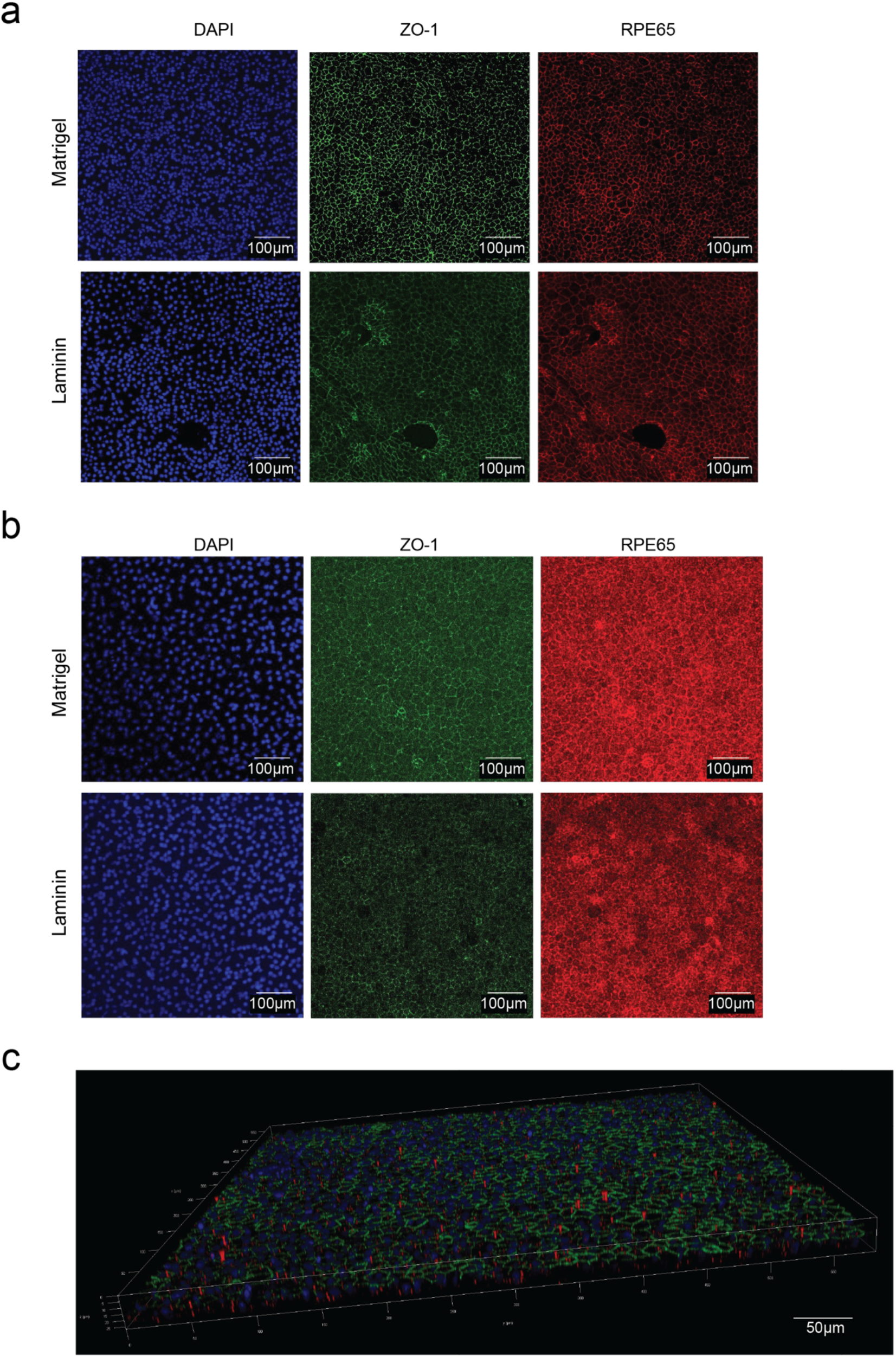
**a**. Differentiated RPE cells in a 2D model with matrigel and laminin. Cells are fluorescently labeled for nucleus (DAPI), ZO-1 (green), and RPE65 (red), **b**. Same as in a. but in a 3D model, **c**. Three-dimensional projection of a 3D model. Cells are fluorescently labeled for nucleus (DAPI), ZO-1 (green), and APOE (red).

**Supplementary Figure 2.**
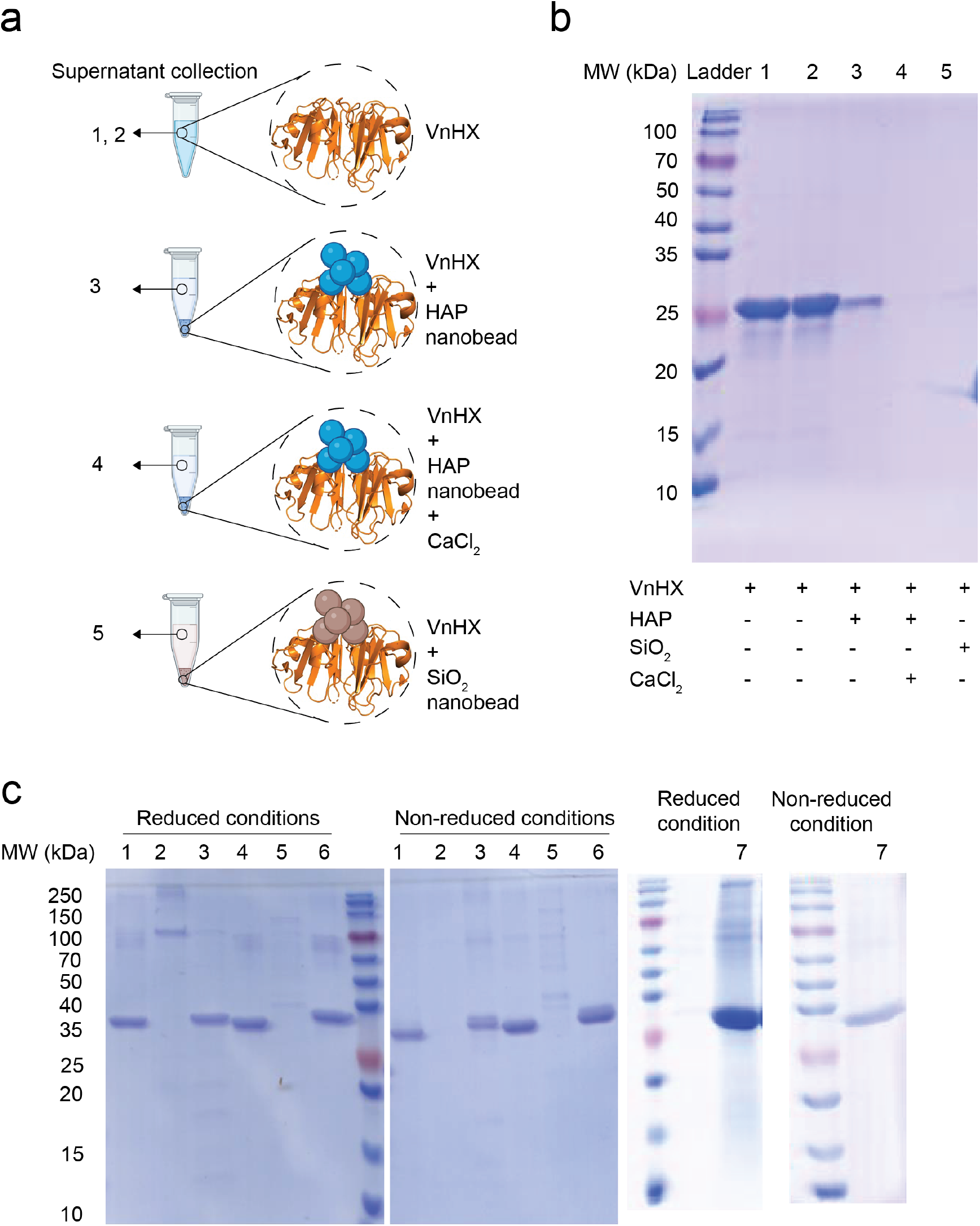
**a**. Schematic of co-sedimentation assay to detect the binding interaction between VnHX and HAP **b**. Reduced PAGE analysis of co-sedimentation assay with Coomassie blue staining for supernatant of a mixture containing VnHX alone (columns 1 and 2), with HAP nanobeads (column 3), with HAP nanobeads supplemented with CaCI_2_ (column 4), and with SiO., nanobeads as a positive control (column 5). **c**. Reduced and non-reduced PAGE analysis of selected scFv expression (lanes 1: NRONC4-4; 2: NRONC4-8; 3: NRONC4-9; 4 NRONC3-7; 5: NRONC3-11; 6: NRONC3-15; 7: NRONC5-1)

**Supplementary Figure 3.**
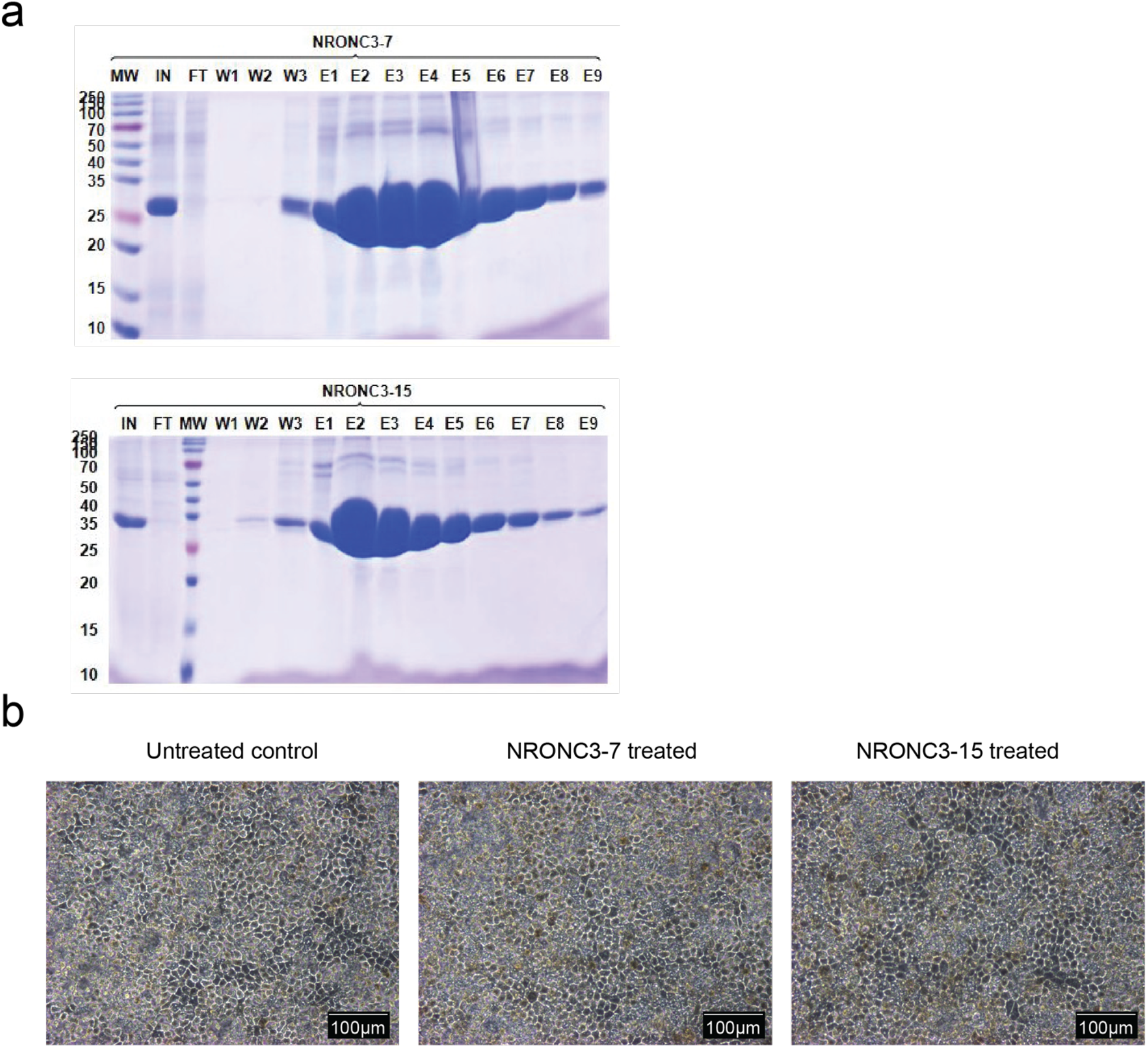
**a**. Purification profile of NRONC3-7 and NRONC3-15 Non-reduced SDS-PAGE analysis conducted with Coomassie blue staining. IN: input sample; FT: flow through: W: wash; E: elution; MW: molecular weight marker (kDa). **b**. Bright field images of RPE in 2D models treated with carrier only (PBS), NRONC3-7 (200 pg/mL), and NRONC3-15 (200 pg/mL), for 48 h.

**Supplementary Figure 4.**
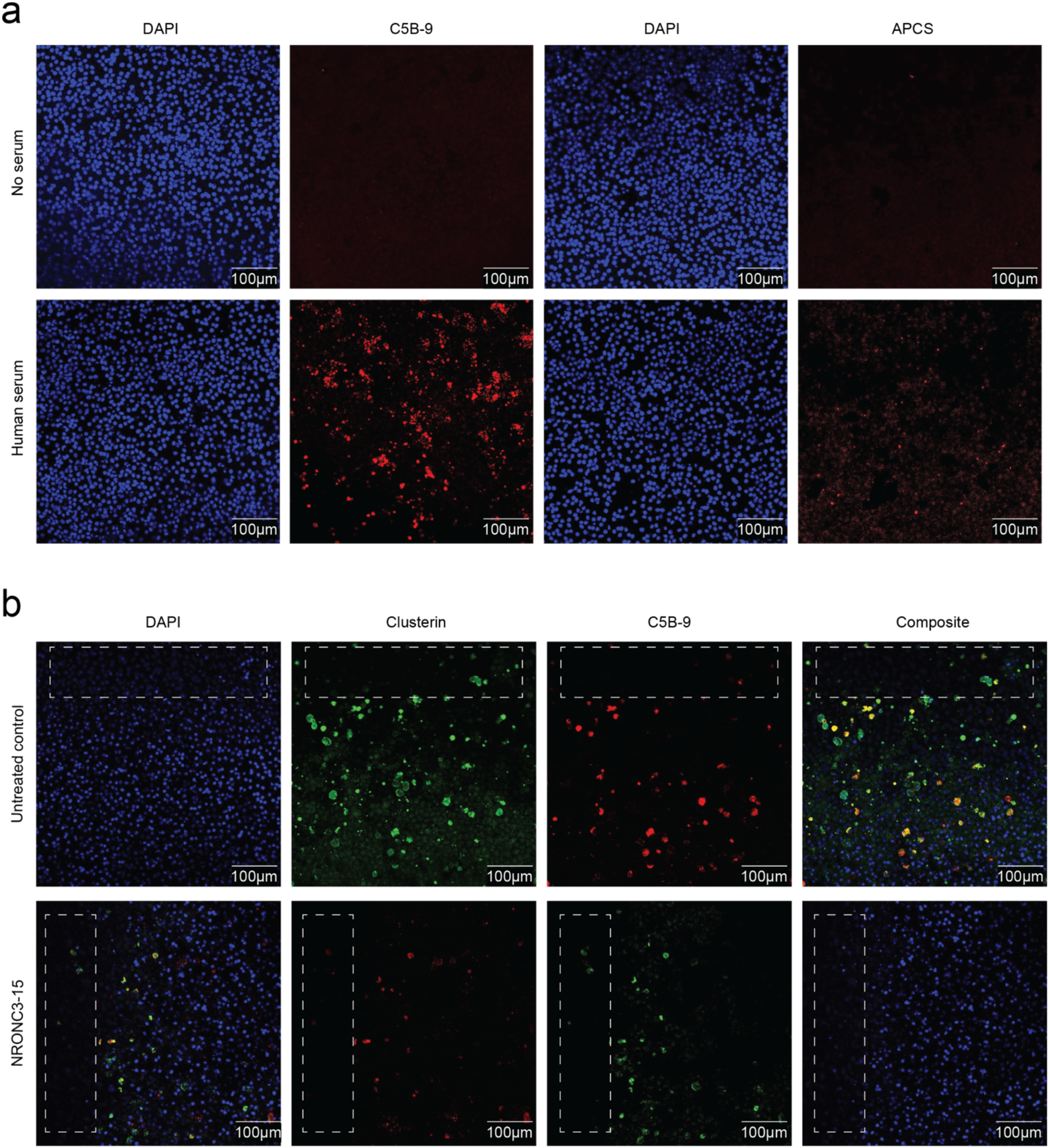
**a.** 2D RPE models on laminin with and without human serum treatment. Models were fluorescently labeled for nucleus (DAPI), C5b-9 (red), and APCS (red), **b.** Expanded images of figure 4d, where control and scFv-treated 3D RPE models on laminin were fluorecently labeled for nucleus (DAPI), clusterin (green), and C5b-9 (red). Areas marked by white dashed lines indicate the membrane bending caused by the long-term culture.

